# Neuronal population correlates of target selection and distractor filtering

**DOI:** 10.1101/422873

**Authors:** Elaine Astrand, Claire Wardak, Suliann Ben Hamed

**Affiliations:** Institut des Sciences Cognitives Marc Jeannerod, Département de Neuroscience Cognitive, CNRS UMR 5229, Université Claude Bernard Lyon I, 67 Boulevard Pinel, 69675 Bron Cedex, France; School of innovation, design, and engineering, Mälardalen University, Högskoleplan 1, 721 23 Västerås, Sweden; Imagerie et Cerveau, INSERM U1253, Université de Tours, Faculté de Médecine, 10 boulevard Tonnellé, 37032 Tours Cedex 1, France

**Keywords:** Frontal Eye Fields, neuronal population, perception, decoding, target selection, distractors

## Abstract

Frontal Eye Field (FEF) single-cell neuronal activity discriminates between relevant and irrelevant visual stimuli and its magnitude has been shown to predict conscious perception. How this is reflected at the population level in terms of spatial codes is unknown. We recorded neuronal population activity in the FEF while monkeys were performing a forced choice cued detection task with identical target and distractor stimuli. Using machine learning techniques, we quantified information about the spatial estimate of targets and distracters in the FEF population activity and we analyzed how these relate to the report of perception. We found that the FEF population activity provides a precise estimate of the spatial location of perception. This estimate doesn’t necessarily match the actual physical world. Importantly, the closer this prefrontal population estimate is to the veridical spatial information, the higher the probability that the stimulus was reported as perceived. This was observed both when the reported stimulus was a target (i.e. correct detection trials) or a distractor (i.e. false alarm trials). Overall, we thus show that how and what we perceive of our environments depends on the precision with which this environment is coded by prefrontal neuronal populations.

## Introduction

Perception is often defined as the ability to become aware of one’s environment through senses. How we perceive our surroundings is influenced both by internal voluntary top-down processes, whereby higher priority is given to relevant aspects of the environment relative to irrelevant aspects, and by external involuntary bottom-up processes, whereby intrinsically salient items impose themselves onto our perception. The outcome of perception can be veridical, reflecting reality, or it can be erroneous, misrepresenting parts of reality or mistakenly interpreting sensory input. In a recent report, Vugt et al. (2018) propose that ignition, i.e. self-sustained prefrontal neuronal activation that bring the neuronal activity to a threshold, accounts for the behavioral report and conscious detection. Weak stimuli that do not evoke neuronal responses beyond early processing levels or transient prefrontal activation do not reach the reportability threshold. In contrast, due to neuronal variability, ignition can be initiated in the absence of a stimulus, leading to false perceptual reports. Pre-existing brain state markers are proposed to have a major effect onto the outcome of perception.

In this respect, the dorsolateral prefrontal cortex, including the frontal eye fields (FEF), is proposed to play a key role in the conscious report of perception (Monosov et al. 2008; Panagiotaropoulos et al. 2012). This is supported by evidence from electrical microstimulation of the macaque FEF leading to enhanced perception (Moore and Fallah 2004) and inactivation of the FEF that induces deficits in visual search target detection independent of search difficulty (Wardak et al. 2006). The neuronal responses of this cortical region to salient unambiguous stimuli are independent of the subject’s overt report (Trageser et al. 2008) and have longer latencies than those observed in the parietal cortex (Ibos et al. 2013). In contrast, the neuronal responses to task-relevant low saliency stimuli emerge in the FEF (Monosov et al. 2008) and precede those observed in the parietal cortex (Ibos et al. 2013). In addition, these neuronal responses highly correlate both with reaction times and response accuracy (Monosov and Thompson 2009) indicating a tight link between FEF activity and overt behavior.

Perception does not only involve selecting a target, but also includes filtering out competing distractors. It has long been established that the activity in FEF demonstrates this ability. Specifically, in pop-out visual search tasks, i.e. in tasks in which the target can easily be distinguished from the distractors, neuronal responses to a distractor presented in the neuron’s receptive field are suppressed (Schall and Hanes 1993; Thompson et al. 1996). In more difficult visual search tasks, as the distracter resembles the target, the neuronal activity in the FEF becomes less reliable at discriminating the target from the distracter (Sato et al. 2003). The degree of distractor suppression as assessed from overt behavior correlates with the degree of neuronal suppression and the inactivation of the prefrontal cortex induces a significant increase in distractibility, i.e. in the production of undesired responses to intervening distractors (Suzuki and Gottlieb 2013).

The prefrontal neural correlates of perception and distractor filtering have most often been studied from the perspective of the response of single neurons. In the present study, we take a neuronal population perspective and apply machine learning techniques to identify how targets and distractors are represented by the FEF population on a given trial and how this representation is predictive of overt behavior. Specifically, we use a forced choice cued target detection task, in which trained monkeys are required to respond as fast as possible to a low saliency target presented at a cued location while at the same time ignoring distractors identical in all respects to the target and presented at other uncued locations. We show that when a stimulus is reported, whether this stimulus is the target of behavior or a distractor, the neuronal FEF population precisely encodes its location. In other words, the population constructs an accurate representation of the stimulus in space. In contrast, when a stimulus is not reported, whether this stimulus is the target of behavior or a distractor, its spatial representation, as encoded by the FEF, does not match its real location. We describe a strong correlation between the error in the estimation of the position of the visual stimulus in space as coded by the neuronal population with respect to its actual physical location, and overt behavior. Overall, we propose that visual perception is not only accounted for by the strength of a visual representation but also by how accurately it is encoded in the neuronal population.

## Methods

### Surgical procedure and FEF mapping

All experimental procedures were identical to those used in Astrand et al. 2016. One head fixation post and two MRI compatible PEEK recording chambers were placed over the FEF one in the left and one in the right hemispheres of two male rhesus monkeys (Macaca mulatta) weighing between 6 to 8kg. During the surgery, gas anesthesia was provided to monkeys using Vet-Flurane, 0.5 – 2% (Isofluranum 100% at 1000 mg/g) followed by an induction with Zolétil 100 (Tiletamine at 50mg/ml, 15mg/kg and Zolazepam, at 50mg/ml, 15mg/kg). Post-surgery pain was controlled with a Morphine pain-killer (Buprecare, buprenorphine at 0.3mg/ml, 0.01mg/kg), 3 injections at 6 hours interval (first injection at the beginning of the surgery) was administered post-surgey and a full antibiotic coverage was provided with Baytril 5% (a long action large spectrum antibiotic, Enrofloxacin 0.5mg/ml) at 2.5mg/kg, one injection during the surgery and thereafter one each day during 10 days. In order to have a precise localization of the arcuate sulcus and surrounding gray matter underneath each of the recording chambers, a 0.6mm isomorphic anatomical MRI scan was acquired post surgically on a 1.5T Siemens Sonata MRI scanner, while a high-contrast oil-filled 1mmx1mm grid was placed in each recording chamber, in the same orientation as the final recording grid. The FEF was defined as the anterior bank of the arcuate sulcus and sites were specifically targeted in which a significant visual and/or oculomotor activity was observed at 10° to 15° of eccentricity from the fixation point during a memory guided saccade task. In order to maximize task-related neuronal information at each of the 24-contacts of the recording probes, we only recorded from sites with task-related activity observed continuously over at least 3 mm of depth. All procedures were approved by the local animal care committee (C2EA42-13-02-0401-01) in compliance with the European Community Council, Directive 2010/63/UE on Animal Care.

### Behavioral task

A 100% validity cued luminance change detection task with temporal distractors (figure 1A) was used. With their head fixed, the monkeys were placed in front of a computer screen (1920×1200 pixels and a refresh rate of 60 Hz). To initiate a trial, they had to hold a bar in front of the animal chair, thus interrupting an infrared beam. On trial initiation, a blue fixation cross (0.7×0.7°) appeared in the center of the screen and the monkeys were required to hold fixation throughout the entire trial, within a fixation window of 2°x 2°. Break of fixation aborted the trial.

**Figure 1.**
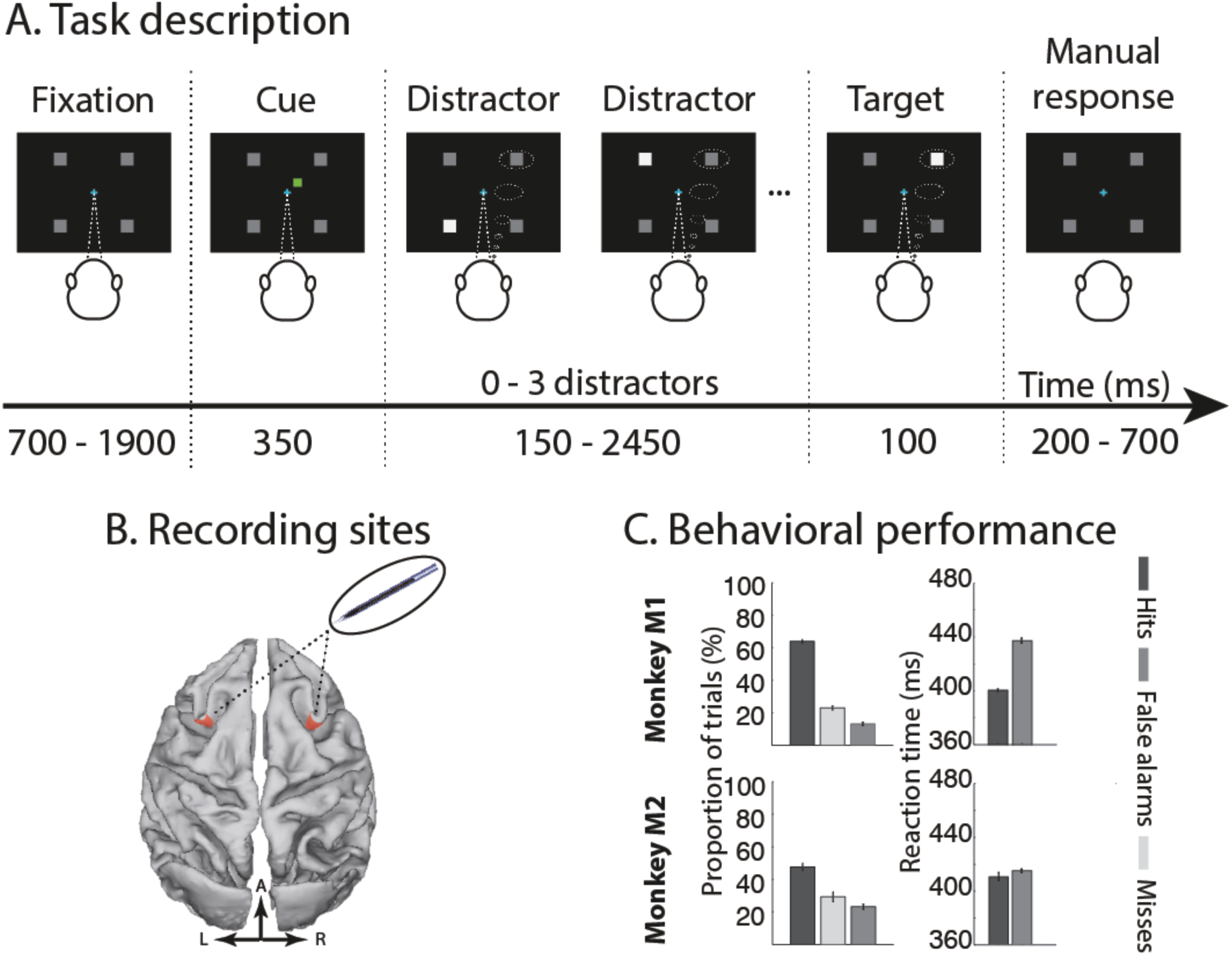
(A) **Task description.** A trial was initiated by the simultaneous onset of a fixation point and four gray landmarks. Monkeys were required to hold a bar and fixate the fixation point throughout the trial. After a variable delay ranging from 700 to 1900ms, a small green square was presented near the fixation point, indicating the location in which the target will be presented. During a variable delay, ranging from 500 to 2800ms, monkeys were required to orient their attention to the cued landmark to detect a small change in luminosity. During the delay, a change in luminosity could occur on any of the other three landmarks and monkeys were required to ignore them. A liquid reward was distributed to the monkeys for releasing the bar 200 to 700ms after target luminosity change. (B) **Recording sites**. On each session, two 24-contact recording probes were placed, one in each FEF. (C) **Behavioral performance**. Median across sessions (n=15) of the proportion of hit trials of (dark gray bars), misses (light gray bars), and false alarms (intermediate gray bars) are depicted for each monkey separately. Error bars correspond to median absolute deviation across sessions.

Four gray square landmarks (0.5°x0.5° for monkey M1, 0.68°x0.68° for monkey M2) were presented simultaneously with the fixation cross and were placed at an equal distance from the fixation point, in the upper right, upper left, lower left and lower right quadrants of the screen, thus defining the corners of an imaginary square. To ensure that the recorded neurons represented the cued spatial location, we adjusted the eccentricity of the landmarks from day to day between 10° to 15°, as inferred from the neurons’ response to a memory-guided saccade task with saccadic targets placed at variable locations in this range. After a variable delay from fixation onset, ranging between 700 and 1900 ms, a green squared cue was presented for 350 ms, indicating to the monkey in which of the four landmarks the rewarding target change in luminosity would take place. The cue was small (0.2°x0.2° for monkey M1 and 0.3°x0.3° for monkey M2) and it was presented close to the fixation cross in the same direction as the landmark to be attended (at 0.3° for monkey M1 and at 1.1° for monkey M2, from the fixation point). After cue presentation, the monkeys needed to orient their attention to the target landmark in order to monitor it for a change in luminosity while maintaining eye fixation onto the central cross. The change in target luminosity could occur anywhere between 500 to 2800 ms from cue onset according to a uniform probability distribution. In order to receive a water or juice reward, the monkeys were required to release the bar (thus restoring the infrared beam) in a time window of 200 to 700 ms following the change in target luminosity (hit trial). In order to make sure that the monkeys were correctly orienting their attention towards the cued landmark, unpredictable changes in the luminosity, identical to the awaited target luminosity change, could take place at the non-cued landmarks (distractors). On each trial, from none to three such unpredictable distractor luminosity changes could take place, no more than one per non-cued landmark position. The monkeys had to ignore these distractors. Responding to such a distractor interrupted the trial and was counted as a false alarm trial if the response fell within 200 to 700 ms following the distractor. Failing to respond to the target (miss trial) similarly aborted the ongoing trial.

### Neural recordings

Bilateral simultaneous recordings in the two FEF hemispheres were carried out using two 24-contact Plexon U-probes. The contacts had an interspacing distance of 250 µm. Neural data was acquired with the Plexon Omniplex^®^ neuronal data acquisition system. The data was amplified 100 times and digitized at 40,000 Hz. The neuronal data was high-pass filtered at 300 Hz. In the present paper, all analyses are performed on the multi-unit activity recorded on each of the 48 recording contacts. A threshold defining the multi-unit activity was applied independently for each recording contact and before the actual task-related recordings started. All further analyses of the data were performed in Matlab.

### Discrete classification procedure

Decoding analyses were performed in Matlab. A regularized linear regression was used to investigate whether the neural population contained information about the variable of interest. A linear regression defines the weight matrix W that minimizes the mean square error of C=W*(R+b), where C is the class (here, the spatial position, amongst four possible locations), b is the bias and R is the neural response (here, a 48 element vector representing the neuronal multi-unit activity at each of the 48 recording contacts, at the time of interest; for each recording channel and each trial). The multi-unit activity was smoothed by averaging the spiking activity over 150 ms sliding windows (resolution of 1 ms); this window width corresponds to a trade-off between decoding performance and decoding speed, as narrower filtering windows result in a lower performance while wider filtering windows decrease temporal resolution (Farbod Kia et al. 2011). To avoid over-fitting we used a Tikhonov regularization which gives us the following minimization equation: norm(**W**\*(**R+b) – C**)+ λ*norm(**W**). The scaling factor λ was chosen to allow for a good compromise between learning and generalization (Astrand et al. 2014). Specifically, the decoder was constructed using two independent regularized linear regressions, one classifying the x-axis (two possible classes: −1 or 1) and one classifying the y-axis (two possible classes: −1 or 1). Each regression used the following procedure. The neural data was divided into training and testing set. The training set was used to define the weight matrix, W and the testing set was then multiplied by W to yield the predicted class C, for these novel trials. Final classification performance was calculated by dividing the number of correct predictions of the classifier on test trials by the overall testing sample size. As a result, the classification performance is a measure, ranging between 0 and 100%, of how much information the neural population contains concerning a specific variable, at a specific time in the trial. When the train- and test-sets of neuronal activities correspond to the same timing relative to the key events of the task, the classification performance is a measure of the instantaneous information content of the variable of interest at exactly that time. When the train- and test-sets of neuronal activities correspond to the different timing relative to the key events of the task, the classification performance is a measure of how the encoding of the encoded information at one moment in the trial generalizes to another moment in the trial.

### Two-tailed non-parametric random permutation

Due to task configuration, absolute chance level is at 25%. However, in order to define the statistical significance of the reported classification performance, we defined for each classifier, the 95% confidence interval limit as follows. For each recording channel, we reassigned random labels (e.g. relative to the cue) to each trial and performed the same classification analysis as described above. This procedure was repeated 1000 times and yielded a 1000 data point distribution of chance classification performance for each combination of a single train time and single test time. Classification performance for real non-permuted data was considered significantly above or below chance if it fell within the 2.5% upper or lower tail of this random permutation distribution (2-tailed non-parametric random permutation test, 0.05 alpha level).

### Continuous classification procedure

As a complement to the above discrete classification procedure, we also investigated the continuous (x,y) output of the classifier that provides a more precise localization of the spatial variable of interest, whether cue representation, spatial attention orientation or perceived target location. The training procedure is identical to that described above in that it trains the classifier to associate the data to the cued (x,y) position, where x and y only takes −1 and 1 values. The testing procedure is slightly different in that it tests novel data without assigning the output of the classifier to a class, but rather to an (x,y) position. The result of this analysis can thus be read as the spatial locus of the variable of interest, whether cue representation, spatial attention orientation or perceived target location, at any point in time. This information can also be represented as a spatial probability map, constructed by calculating the statistical significance of a given (x,y) classifier output at each spatial location using non-parametric random permutation test (see previous section) and visualizing these as a color coded z-scores. Specifically, a map of p-values was constructed by comparing the actual data with randomly permuted data. Z-scores were then calculated from these p-values yielding the signed number of standard deviations from the normal mean probability, at each spatial location, that is significantly over- or under-represented (p<0.01).

### Euclidian Multidimensional distance

The multidimensional Euclidian distance (hereon named MDD) between multiunit activity for all stimulus position-pairs was calculated per trial as follows: 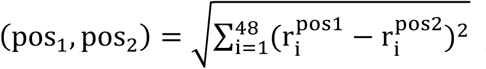, where r corresponds to the neuronal response for each channel i.

The MDD for all position-pairs were averaged to yield one MDD-value per trial. Trials were binned in 20 equally sized bins according to the distance between target location for hit trials (resp. distractor location, T(D), that elicited the response for false alarms) and the decoded target location (resp., T(D), for false alarms). Corresponding MDD-values were averaged.

## Results

### Behavioral performance

Monkeys performed a spatially cued target detection task. Trials were divided according to their overt behavior as follows. A trial was considered correct (hit trial) if a response was produced between 200ms and 700ms from the onset of target luminance change. A trial in which the monkeys did not produce any response to a target was considered a miss. If a response was produced 200ms to 700ms following a distracter luminance change, this trial was considered a false alarm trial. M1 and M2 achieved 64.4% and 47.9% correct responses, respectively (figure 1C). Reaction times on these correct trials were on average 401ms and 410ms, respectively. Both monkeys tended to produce more misses than false alarms (M1, Miss = 22.5%, FA = 12.9%; M2, Miss = 27.9%, FA = 23.0%, p<0.001, Wilcoxon paired test) and reaction times were notably longer on false alarm trials compared to correct trials (p<0.01 for both monkeys, Wilcoxon paired test).

### Spatial attentional prioritization of the target

In the following, we quantify the available information about attention orientation in the prefrontal FEF neuronal ensembles being recorded from, as a function of the overt behavior of the monkeys. Specifically, recorded multi-unit activity (MUA), on correct trials, was used to train two regularized linear regressions (one along the x-axis and one along the y-axis) to associate bilateral neural response patterns at a given time from target presentation with the location of expected target, and this for successive time windows around target presentation. Figure 2A shows the resulting instantaneous classification performance, in time, around target presentation across all recording sessions. This classification performance represents the percentage of trials for which the classifier assigned its output to the cued quadrant. Confirming our previous report (Astrand et al. 2016), during correct trials (figure 2A, grey) before target presentation, classification performance represents attentional prioritization at the cued location. Accuracy was around 40% and was maintained above the 95% confidence interval limit (2-tailed non-parametric random permutation), indicating that the monkeys were actively orienting their attention to the cued spatial location. In contrast, for miss trials (figure 2A, pink) prior to target presentation, classification performance was around chance (25%) suggesting that, in these trials, the monkeys did not consistently orient their attention to the cued location.

**Figure 2.**
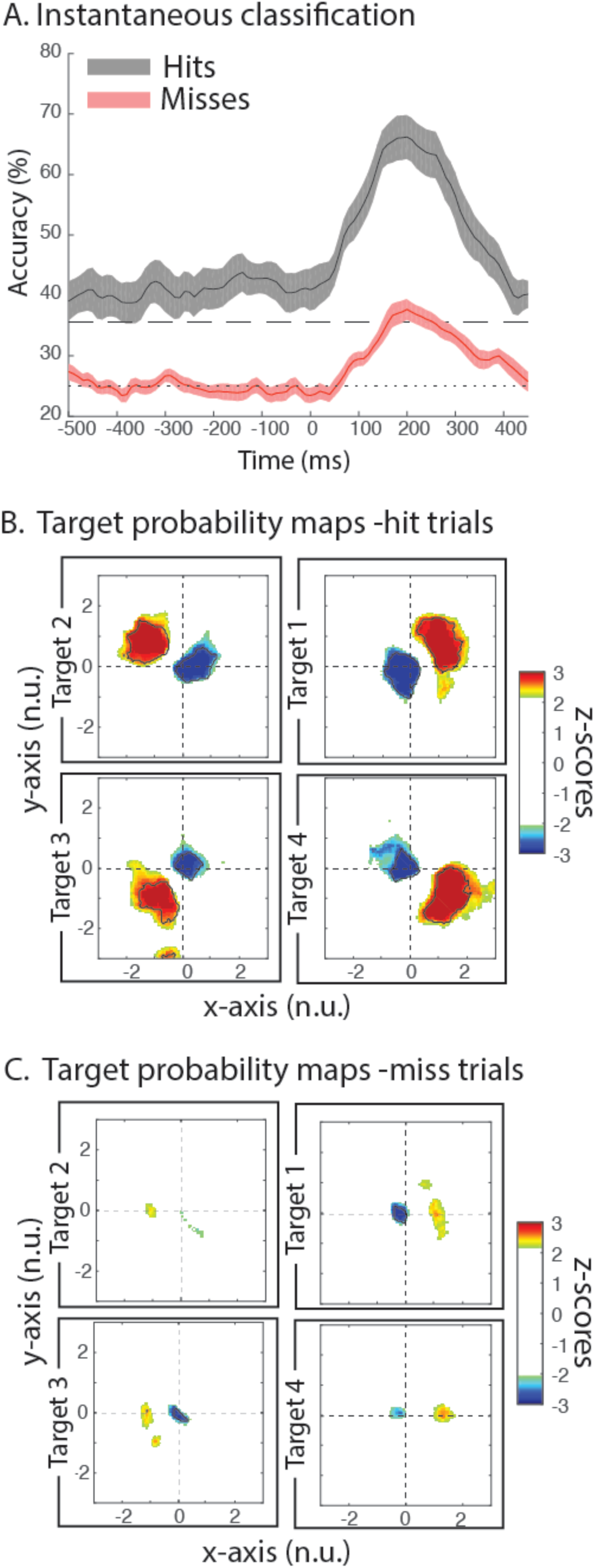
Target perception. (A) Instantaneous classification accuracy of target location in time, aligned on target presentation (time=0ms). Graphs show mean classification accuracy (%) with associated standard error across sessions (n=15) for hits (gray) and miss trials (red). At each time point the train-time of the classifier is the same as the test-time. The dotted horizontal line corresponds to chance classification performance (25%) and the striped horizontal line corresponds to the upper 95%-confidence interval determined by random permutation tests. (B-C) Target (x,y)-location probability maps for hit trials (B) and miss trials (C). Maps represent z-scores of the spatial locations that are statistically over-represented (red color-scale, p<0.01) or under-represented (blue color-scale, p<0.01) with respect to the 95%-confidence interval (determined by random permutation tests) when decoding target location 100 to 200ms after target presentation, over all sessions and both monkeys. Locations (10° or 13° eccentricity depending on the session) are normalized across sessions so that target locations are mapped to an eccentricity of +/- 1. Each map corresponds to one of the four cued locations (e.g. upper right quadrant representing the map of spatial perception distribution for target appearing in the upper right quadrant). The black contour indicates the conjunction of mean z-scores over all sessions and both monkeys.

### Target perception

Instantaneous classification performance after target presentation (Figure 2A) reflects how much information is available in the FEF population relative to target location. On correct trials, target-related information peaks at 200ms, with an average of 65% correct predictions. This increase is well beyond the 95% confidence interval, and statistically significantly higher than the pre-cue attentional orientation related signals (comparing accuracy 100ms to 200ms post-target onset, with accuracy 200ms to 100ms pre-target onset, average difference, 28%, p<0.001, Wilcoxon paired test). On miss trials, significant increase across sessions is also observed although it barely reaches the upper 95% confidence interval limit (average difference 9%, p=<0.001, Wilcoxon paired test). This is in agreement with the findings that attentional orientation towards the receptive field of FEF neurons facilitates the detection or the discrimination of a target, showing a lower spiking rate on miss trials than on correct trials, both in the attention orientation period and following target presentation (Thompson and Schall 1999; Ibos et al. 2013).

Whenever an item is perceived in our environment, it is implicitly associated to a more or less precise location in space. To investigate this aspect of target perception, the continuous output from the classifier was used to predict the spatial location of perception using the same procedure that we have previously reported to track the attentional locus from FEF neuronal activities (Astrand et al. 2016). In the following, we focus on a time-period running from 100ms to 200ms after target presentation. Figure 2B and 2C represent the (x,y) locations decoded from the population activities that were statistically overrepresented (red color scale, z-scores > 2.33, p<0.01) or underrepresented (blue color scale, z-scores < −2.33, p<0.01) as a function of the position of the target with respect to chance (estimated by a 2-tailed non-parametric random permutation test). On correct trials (Figure 2B), a rather large area of overrepresented decoded locations around the location at which the target was actually presented can be observed for all four target locations (hot colors, figure 2B, average over all sessions, contour shows conjunction between all sessions and both monkeys). In contrast, a smaller area of underrepresented decoded locations can be observed in proximity to the fixation point for each target location (cold colors, figures 2B). In a subsequent analysis, we examine the relation between attentional locus and spatial location of perception as estimated from the prefrontal neuronal population. We observe a strong correlation between the two processes. In correct trials during which attention was oriented close to the cued location prior to target presentation (attention located within 7° of the cued location and the fixation point 300ms to 200ms prior to target presentation), the perceived location of the target (100ms to 200ms after target presentation) was significantly closer to its real physical location compared to trials during which attention was oriented outside the quadrant of the cued location prior to target presentation (close attention trials: distance = 8.8 +/- 2.2, far attention trials: distance = 11.9 +/- 1.9, p=0.00036). This indicates a strong relationship between preorientation of attention and the location of perception. In other words, the precision with which attention is preoriented to the cued location will strongly influence the precision of the perceived spatial location of the target.

On miss trials (Figure 2C, decoder trained on hit trials and tested on miss trials), only weakly overrepresented decoded locations can be identified. These locations fall short off target location and there is no conjunction between all sessions and the two monkeys. Underrepresented decoded locations can be identified around the fixation location, though they are smaller than those identified on correct trials (figure 2B). Overall, this indicates that correct target detection correlates with a reliable decoder localization of the target in the vicinity of its actual location. In contrast, missed targets correlate with an unreliable localization of the target. Overt perception and precise spatial information thus seem to be tightly related. To confirm this, we further probe this relationship between overt perception and precise spatial information on false perception trials, i.e., false alarm trials.

### Spatial attentional prioritization of the distracter

To investigate how the distracters, presented at a random time during the delay between cue and target presentation, were perceived in the different types of trials, the regularized linear regression was trained to associate MUA with the location of the target and tested onto distracter localization. Figure 3A shows the instantaneous classification performance in time around distracter onset. On false alarm trials (figure 3A, purple), spatial information about the upcoming position of the distractor is significantly above the upper 95% confidence interval limit (two-tailed non-parametric random permutation). Because distractor location was completely pseudo-randomized across trials, there was no way for the monkeys to predict neither distractor probability of presentation, nor distractor time of presentation, nor distractor location. Thus, this spatial bias rather indicates that false alarm trials arise from misorientation of attention at the location before and at the time of presentation of the target (see Astrand et al., 2016 for a detailed analysis of this aspect). On correct (figure 3A, grey) and miss trials (figure 3A, pink), classification performance is well below chance (25%) almost reaching the lower 95% confidence interval limit. This indicates that, on these trials, attention is oriented away from the location of the upcoming distracter –thus accounting for it not being perceived on these trials. Thus, overall, the coincidence between distractor location and pre-distractor spatial attentional priority information contribute to overt behavior. As described below, this is mediated by an enhanced representation of the distractor when it is selected by attention.

**Figure 3.**
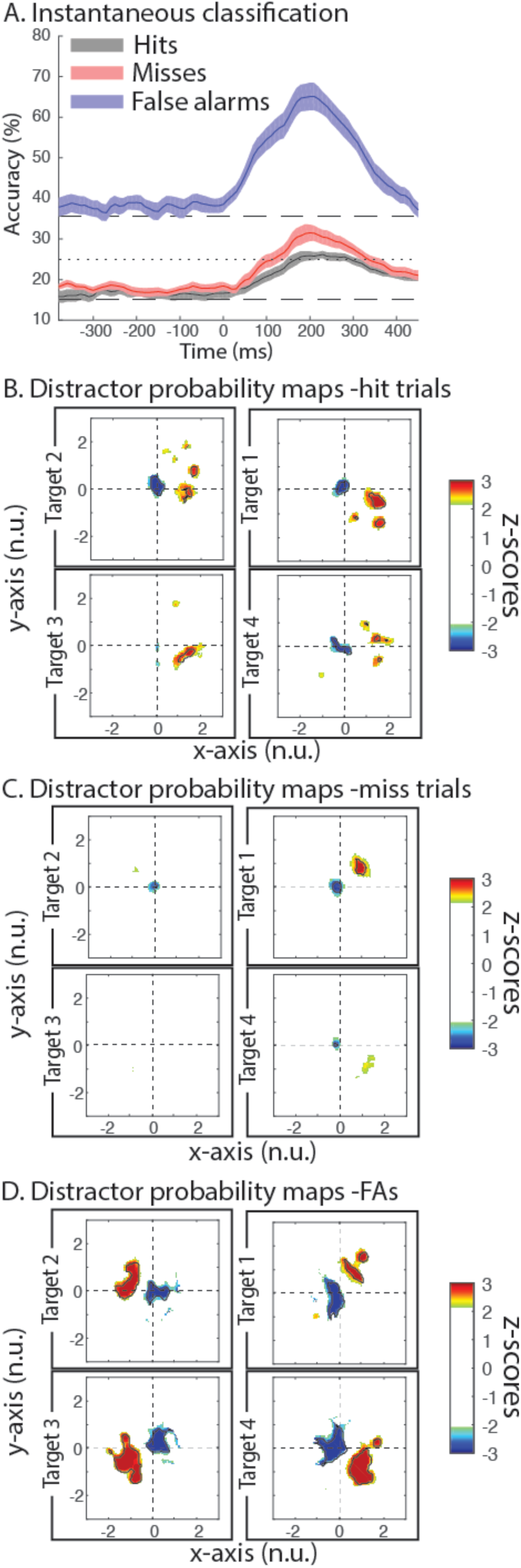
Distracter perception. (A) Instantaneous classification accuracy of distracter location in time, aligned on distractor presentation (time=0ms). (B-D) Distractor (x,y)-location probability maps for hit trials (B), miss trials (C) and false alarm trials (D). All as in figure 2.

### Distractor perception

Following distracter presentation, classification performance increases substantially during false alarm trials (purple) and reaches the same accuracy as for correct trials following target onset (correct trials, average accuracy from 100 to 200ms post target onset: 61%, false alarm trials: 60%, p=0.64, Wilcoxon paired test). On correct (gray) and miss (pink) trials, there is a substantially smaller increase in decoding accuracy following distracter presentation that remains below the upper 95% confidence interval limit (correct trials: average difference 7% below the 95% c.i., p=<0.001, miss trials: 11% below the 95% c.i., p<0.001). In other words, as seen for target perception, distractor perception (as defined by overt false alarm behavior) coincides with a high decoding accuracy of distractor quadrant location in visual space. In contrast, absence of distractor perception coincides with a very low decoding accuracy.

Precise distractor localization in visual space also depends on whether the distractor has been perceived or not. Figure 3B-D represent the (x,y) locations decoded from the population activities that were statistically overrepresented (red color scale, z-scores > 2.33, p<0.01) or underrepresented (blue color scale, z-scores < −2.33, p<0.01) as a function of the position of the distractor with respect to chance (estimated by a 2-tailed non-parametric random permutation test). During both correct (figure 3B) and miss (figure 3C) trials, distractor location is either weakly or erroneously represented. In contrast, on false alarm trials (figure 3D), i.e. trials in which the distractor was mistakenly reported as a target, distractor location is significantly reported at the actual location of the distractor, very much like what is reported for target localization on correct trials in figure 2B. Overall, this thus indicates that stimulus perception (whether a target or a distractor) correlates with a reliable stimulus localization that resides in the proximity of the real stimuli position.

### Distracter vs. target prefrontal representations

By task design, distractors and targets correspond to the same visual stimulus. What distinguishes one from the other is whether the spatial location where the stimulus is presented has been cued or not. In the following, we quantify the similarity between how the FEF encodes, at the population level, target and distractor spatial information. To this goal, a regularized linear regression was trained on correct trials, to discriminate between target and distracter, irrespective of stimulus position. Training was performed on MUA neuronal responses 100ms to 200ms following target or distracter presentation. Figure 4A shows decoding accuracies on novel correct (black), miss (pink) and false alarm (purple) test trials. Specifically, on correct trials, the classifier succeeded in discriminating the target from the distracter in 71% of instances (chance at 50%). On false alarm trials, decoding accuracies were in the same range as those observed on correct trials, and significantly above chance (69%, p=0.12, Wilcoxon paired test, note that the distracter that evoked the response was considered to be the target in these trials). In contrast, on miss trials, decoding accuracy was significantly lower and hardly above chance (54%, p<0.001, Wilcoxon paired test). Overall, this thus suggests that while information about the selected item is well represented in the FEF (target on hit trials and distractor on false alarm trials), hardly any information is available about the unselected item (target on miss trials).

**Figure 4.**
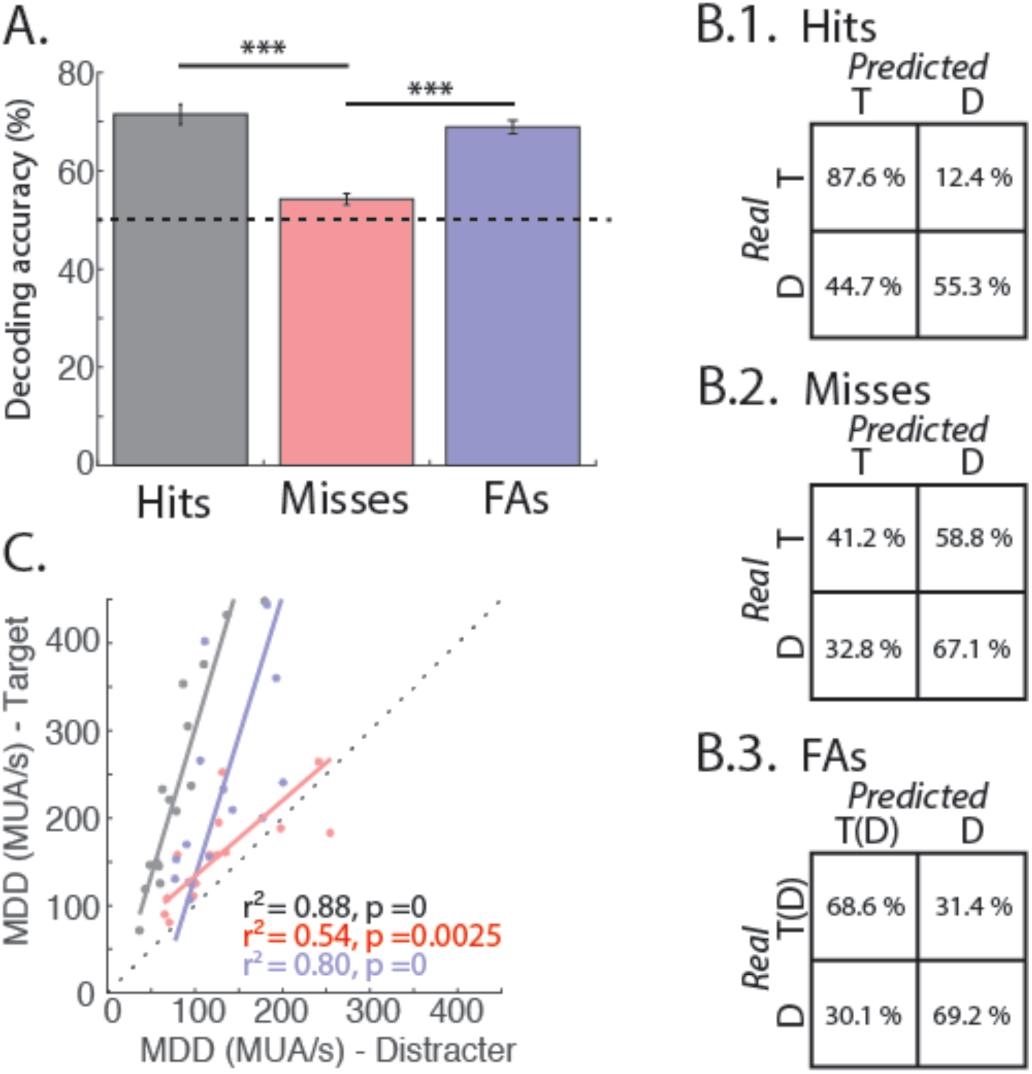
Target and distractor related information. (A) Decoding accuracy at classifying target vs. distractor (mean +/- s.e., in %). Classifier is trained on hit trials and tested on hits (gray), misses (pink) and false alarm trials (purple). Accuracies are calculated over a 100 to 200 ms post-target or post-distractor time interval. Dotted horizontal line corresponds to chance level (50%). (B) Confusion matrices of the classification in A for hit trials (B.1), misses (B.2), and false alarms (B.3). Rows correspond to actual presented stimulus (distracter or target) and columns correspond to the predicted stimulus by the classifier. For false alarms, the target is taken as distractor that elicited the response T(D), all other stimuli are considered as distractors. (C) Euclidian multidimensional distance of MUA over all channels: target related response (y-axis) vs. distracter related response (x-axis) for hit (gray), miss (pink), and false alarm (purple) trials. Each dot corresponds to one session. Solid lines correspond to an orthogonal regression fit. Corresponding Spearman’s correlation statistics are indicated.

The analysis of the confusion matrices refines this view (figure 4B). On hit trials (figure 4.B1), while the target is correctly decoded as a target in 88% of instances, distractors are correctly decoded as distractors in only 55% of instances (88% vs 55%, p<0.001, Wilcoxon paired test). In other words, the distractor is often mistaken for a target in 45% of instances. On miss trials (figure 4.B2), this pattern is reversed. Namely, the target is correctly detected as a target in only 41% of the trials (to be compared to the 88% correct classification on hit trials), while distractors are correctly decoded as distractors in up to 67% of instances. On false alarm trials (figure 4.B3), both the decoded distractor (named T(D), due to the fact that it is selected as a target by the monkey) and the ignored distractor (D) are both correctly classified as target (69%) and distractor (69%) respectively. This suggests that, coexisting with the stronger selection process on hit and false alarm trials, described in the previous paragraph, an overall weaker distractor filtering process is at play during miss and false alarm trials, as available information about distractors is much higher on these trials than on hit trials. This possibly relates to the higher noise correlation observed on miss and false alarm trials relative to hit trials that we previously described in the FEF neuronal population (Astrand et al. 2016).

In order to gain a better understanding of how the representations of a given visual item, as assessed from the neuronal population response patterns, vary as a function of overt behavior, we proceeded as follows. We computed the Euclidian distance in the multidimensional neuronal space (hereon simply called multidimensional distance, MDD) between the neuronal response patterns of the recorded populations to the four possible stimuli locations for two conditions (see section method). We first computed the MDD-target (figure 4.C, y-axis), when the presented stimulus was an expected target (hits and misses) or a distractor that elicited a response (false alarms).

We then computed the MDD-distractor (figure 4.C, x-axis), when the presented stimulus was a non-selected distractor stimulus. For both computations, distances were computed using neuronal activities averaged over 100 to 200ms post-stimulus presentation. MDD following target and distracter presentations are strongly correlated for all three types of trials (figure 4C, hits, gray: p<0.001, r^2^= 0.88; misses, pink: p<0.001, r^2^=0.80; false alarms, purple: p<0.01, r^2^=0.54). Furthermore, MDD is significantly higher following target presentation as compared to after distracter presentation for correct trials (average 238 MUA/s vs. 81 MUA/s, p<0.001, Wilcoxon paired test). The same relationship can be observed for false alarm trials when considering the distracter that evoked the erroneous response as the target (average 244 MUA/s vs. 125 MUA/s, p<0.001, Wilcoxon paired test). During miss trials there is a moderate though significant increase in the MDD following target presentation as compared to after distracter presentation (157 MUA/s vs. 127 MUA/s, p<0.01). This may indicate the co-existence of an active selection process associated with an overt behavioral response (in hits and false alarm trials) which coincides with enhanced neuronal responses following the visual stimulus, and an active suppression mechanism that is associated with the absence of overt behavioral response which coincides with decreased neuronal response following a visual stimulus (i.e. compare neuronal responses following a distractor in hit trials with those following a target in miss trials).

In a last step, we sought to test whether these active filtering and selection processes were associated with a change in the coefficient of variation of the neuronal responses. We thus calculated for each channel, on each session, the ratio between the standard deviation and the mean, following target or distractor presentation, for each trial type (hits, misses and false alarms). A 2-way ANOVA (trial type x target vs. distracter) revealed a significant main effect of target vs. distracter presentation (p<0.001). Further post-hoc tests show that this coefficient of variation was significantly lower following target as compared to after distracter presentation for correct trials (target: 0.30, distracter: 0.32, p<0.001, Wilcoxon paired test). This was also true during false alarm trials (distractor eliciting the response: 0.30, distracter: 0.31, p<0.05). During miss trials, no difference was observed (target: 0.31, distracter: 0.32, p=0.42). In other words, stimulus selection was associated with more reliable neuronal responses and a 10 to 20% decrease in the coefficient of variation of the neuronal responses.

### Behavioral correlates of perceived target/distractor location

In Astrand et al. (2016), we show that the distance between the attentional spotlight, as decoded from FEF activity account for overt behavior: 1) the closer the spotlight from the upcoming target, the lowest the miss rates and the fastest the reaction times; 2) the closest the spotlight from the upcoming distractor, the highest the false alarm rates and the fastest the reaction times. The same holds true for the distance between the perceived location of the target or distractor, as decoded from FEF activity (figure 5). Specifically, both monkeys show a significant correlation between median reaction times and mean distances between actual target location and (x,y) decoded target location in 20 equally sized distance bins (estimated on average neuronal responses from 100 to 200ms after target onset), on correct trials (figure 5A, M1: r^2^ = 0.42, p<0.001, M2: r^2^ = 0.35, p<0.001). Likewise, a significant correlation is observed on false alarm trials between median reaction times and mean distances between actual distracter location (the distractor that evoked the response) and (x,y) distractor decoded location, in 20 equally sized distance bins, estimated on average neuronal responses from 100 to 200ms after distractor onset (figure 5B, M1: r^2^ = 0.16, p<0.05, M2: r^2^ = 0.40, p<0.001). In addition, the proportion of misses over correct trials increases as the distance between target location and (x,y) target decoded location increases (figure 5C, M1: r^2^ = 0.79, p<0.001, M2: r^2^ = 0.79, p<0.001). In accordance, as the distance between distracter location (the distractor that evoked the response) and (x,y) distractor decoded location decreases, the proportion of false alarms over correct trials increases (figure 5D, M1: r^2^ = 0.77, p<0.001, M2: r^2^ = 0.93, p<0.001). These results provide a strong indication that where visual items are perceived in space, whether targets or distractors, influence both reaction times and overt responses.

**Figure 5.**
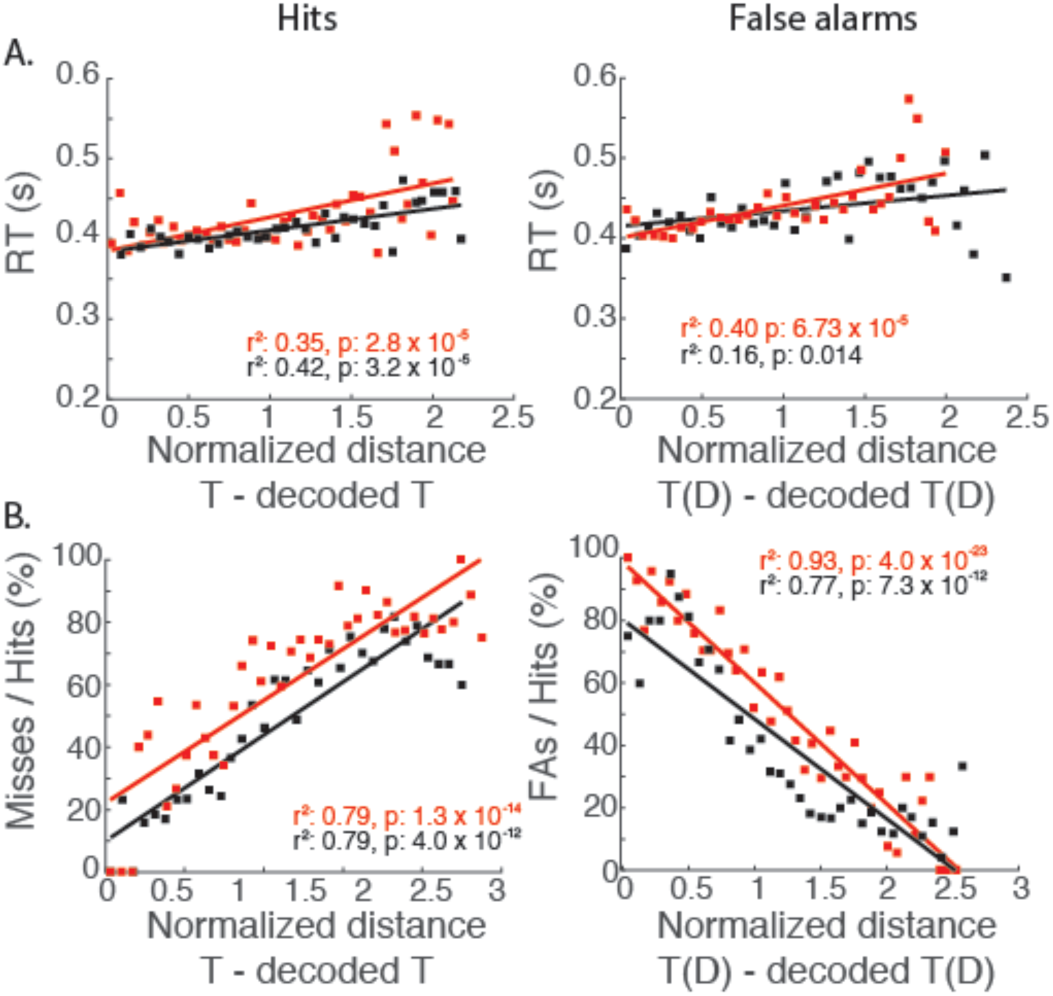
Reaction times (A) and detection performance (B) as a function of target to decoded target distance. Target to decoded target distance are calculated on a time interval running from 100 to 200ms post target presentation. For false alarms, the target is taken as the distractor that elicited the response T(D). Data are represented for hits (left panels), and false alarms (right panels). Each dot corresponds to the mean distance and median reaction times (A) or mean trial-type proportion rate (B) in each out of 20 equally sized distance bins (black, monkey M1, red, monkey M2). Data was fitted with an orthogonal regression (solid lines) and the corresponding statics of Spearman’s correlation are indicated. Locations (10° or 13° eccentricity depending on the session) are normalized across sessions so that target locations are mapped to an eccentricity of +/- 1.

### Population target location estimates reflect onto underlying MUA

The distance between the actual visual stimuli and where they are represented in space as decoded from the FEF neuronal responses is a population estimate. Here, we probe how much this population estimate reflects onto the underlying MUA responses. In other words, do MUA responses correlate with decoder output? Figure 6 represents the correlation between the normalized amplitude of MUA and the distance between target (distracter evoking a response for false alarm trials) location and (x,y) decoder output. MUA signals that had a significant modulation of their response, on correct trials, in the 100 to 200ms time window following target presentation, as compared to the −200 to −100ms prior to target presentation (Wilcoxon paired tests, p<0.05), were selected for further analysis. The target-to-decoder output distance was averaged within 20 equally sized distance bins and the MUA from the corresponding trials within each bin was averaged. On correct trials (figure 6, gray), a significant correlation between target-decoder output distance and MUA amplitude can be observed (r^2^ = 0.48, p<0.01). On false alarm trials, overall MUA amplitude is lower compared to hit trials (p<0.001, Wilcoxon test) but a significant correlation can also be observed between MUA amplitude and distracter-decoder output distance (r^2^ = 0.32, p=0.01). On miss trials, the MUA amplitude is substantially lower as compared to hit trials (485 MUA/s vs. 360 MUA/s, p<0.001, Wilcoxon test). For these trials, a trend towards significance can be observed between the correlation between MUA amplitude and target-decoder output distance (r^2^ = 0.23, p=0.06). Overall, FEF MUA activity thus reflects perceived location, rather than actual stimulus physical location.

**Figure 6.**
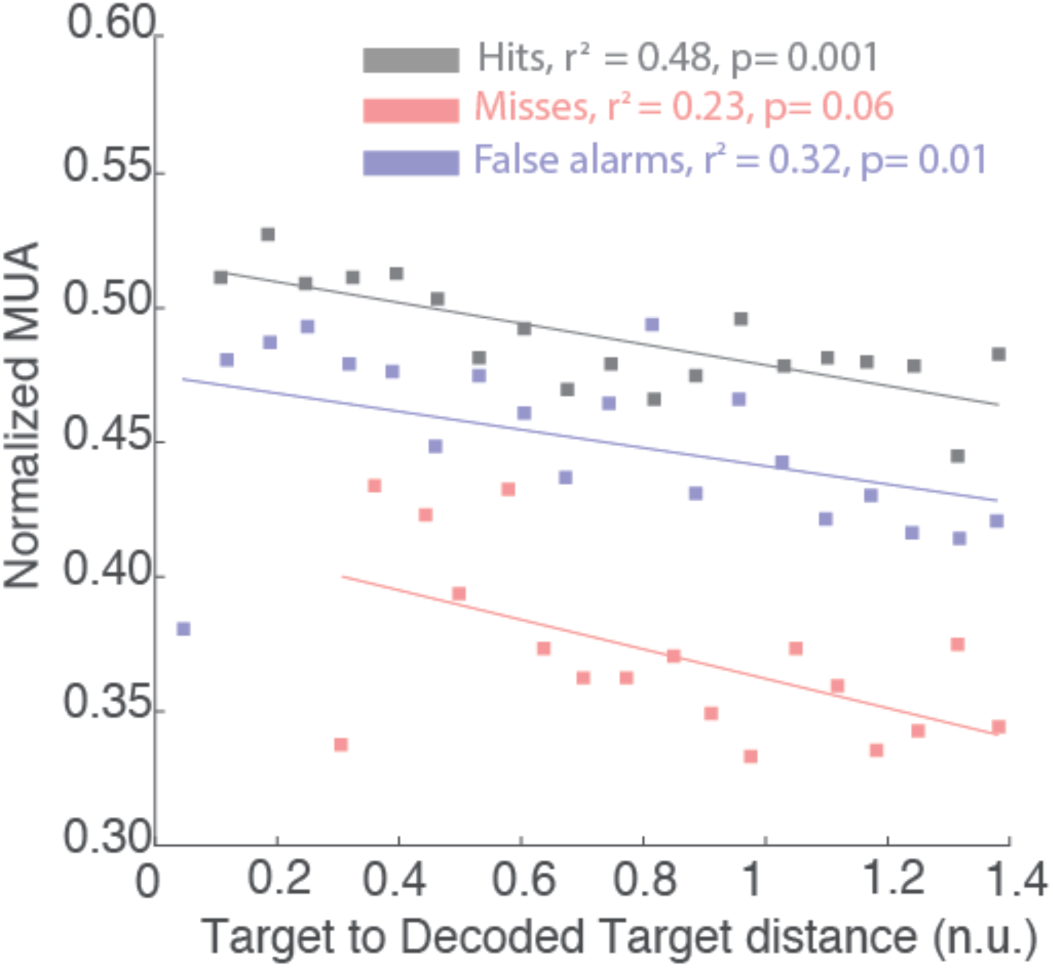
Normalized MUA as a function of target to decoded target distance. Normalized MUA and target to decoded target distance are calculated on a time interval running from 100 to 200ms post target presentation. Data cumulated for both monkeys are represented for hits (gray), misses (pink) and false alarms (purple). In the case of false alarms, measures are extracted relative to distractor presentation. All else as in figure 5.

## Discussion

Visual perception is defined as the conscious representation of a visual item. It is experimentally assessed by requesting an overt report by the subject that can take different forms: a detection, a discrimination, a verbal report etc. In this work, we identify the neuronal prefrontal correlates of visual perception from a neuronal population perspective (Astrand et al. 2014, 2015, 2016). Specifically, we show that when a stimulus is reported, whether this stimulus is the target of behavior or a distractor, the neuronal FEF population precisely encodes its location. In contrast, when a stimulus is not reported, whether this stimulus is the target of behavior or a distractor, its location, as encoded by the FEF, doesn’t match its actual location. We describe a strong correlation between the error in the estimation of the position of the visual stimulus in space as coded by the neuronal population with respect to its actual physical location, and overt behavior. These observations are discussed below.

### Visual perception

In the absence of attentional pre-orientation (cued), single-unit neuronal activity in the FEF during pop-out visual search, driven by bottom-up mechanisms, has been shown to reliably encode the presence of a visual stimulus within the neuron’s receptive field (RF) independently of whether the stimulus is a target or a distracter (Trageser et al. 2008). This indicates that, in this task, the FEF does not contribute to the behavioral report. In contrast, on a difficult cued target detection task, driven by top-down mechanisms, the speed and accuracy of the behavioral response on individual trials is predicted by the magnitude of single neuron responses to the target when presented in the neuron’s RF, this magnitude being lower both on error trials and on trials with longer response times (Monosov and Thompson 2009). The spatial selection of this non-salient task-relevant visual information (target) arise in the spiking activity prior to the local field potentials of the FEF, suggesting that spatial selection emerges in the prefrontal cortex (Monosov et al. 2008). Corroborating this observation, the neuronal responses to non-salient targets arise earlier in the prefrontal cortex than in the parietal cortex (Ibos et al. 2013). At the neuronal population level, we show very high classification rates of both targets and distractors when these are selected to produce a behavioral response (hits and false alarm trials). This translates into a confined localization on the decoding probability maps around the actual physical location of the target/distractor. In other words, on these trials, the location of the target or of the distractor is precisely represented in the prefrontal cortex. In contrast, classification rates of both targets and distractors are below the 95% confidence interval when these are not selected (misses and correct rejections). This corresponds to an unreliable localization on the target/distractor decoding probability maps. In other words, visual stimulus selection correlates with a reliable neuronal population representation of stimulus location on a trial-to-trial basis.

### Prefrontal sensory representations accounts for perception and overt behavior

As discussed in the previous section, in cued target detection tasks, the speed and accuracy of the overt response on individual trials is predicted by the magnitude of single neuron responses to a target in the RF (Monosov and Thompson 2009). Here we show that the spatial estimate of target/distractor location provided by the FEF neuronal population accounts for behavior. Specifically, an accurate spatial representation coincides with 1) shorter reaction times, 2) lower proportions of misses and 3) higher proportions of false alarms when the location of the target/distractor is estimated at its veridical position as compared to further away (figure 5). In other words, the population estimate of target/distractor location parametrically accounts for behavior, with an explained variance of miss and false alarm rates ranging between 70% and 90%. Multi-unit activity in response to the target/distractor presentation also co-varies with this population estimate. However, in this case, the explained variance is much lower and ranges between 20% (MUA response to target in misses) and 50% (MUA response to target in hits). This indicates that the neuronal population better accounts for overt behavior than single neuron or multi-unit activity.

### Target selection vs. distractor filtering

Suzuki and Gottlieb (2013) show that the dorsolateral prefrontal cortex single-unit neural spiking activity following a distracter (that is identical to the target) is positively correlated to error rates. On a population level we corroborate and extend this finding by showing that behavioral performance, in terms of accuracy and response times, correlates with the spatial perceptual representation of a distracter in the FEF. Specifically, we show that as the distance between the FEF neuronal population estimate of the locus of perception and the actual position of the distracter decreases (i.e. as the error of the spatial estimation decreases), the false alarm rate increases and response times during false alarm trials decrease. We further observe that multiunit activity following distractor presentation negatively correlates with the error between the perceived distracter location in false alarm trials and its actual physical location. This indicates that the selection of a distractor for behavioral report co-varies with this perceptual distance error measure. The shorter the distance the higher the selection. Likewise, target selection co-varies in a similar manner with the error between the perceived target location in hit trials and its actual physical location. We similarly observe that the multiunit activity following target presentation negatively correlates with this distance error measure.

Hit trials correspond to trials in which the target has been selected. In contrast, false alarm trials correspond to trials in which a distractor failed to be filtered. By task design, target and distractors were identical physical stimuli, the only distinguishing factor being whether the stimulus was presented at the cued location or not. As a result, one expects that, when perceived, targets and distractors would be represented in identical manners. Average normalized multi-unit in response to a detected target or to a detected distractor were significantly higher than the average normalized multi-unit in response to a missed target. Average normalized multi-unit in response to a detected target was only slightly stronger than the average normalized multi-unit in response to a detected distractor. The neuronal population doesn’t discriminate between an unselected target and an unselected distractor. Likewise, the neuronal population equally discriminates between selected and unselected target and selected and unselected distractors. Overall, this suggests that once a stimulus has been selected, targets and distractors are undistinguishable to the neuronal population. But what triggers selection?

### Interactions between attention and perception

Under low signal or high noise conditions, spatial attention has been shown to facilitate perception at the locus of attention. Indeed, behavioral responses to attended stimuli are faster (Yantis and Jonides 1990) and visual sensitivity at attended locations is enhanced (Bashinski and Bacharach 1980; Carrasco 2011). At the neuronal level, attention has been proposed to operate through a variety of mechanisms including enhanced neuronal response to visual stimuli when attention is oriented towards the receptive field of the neuron (e.g. McAdams and Maunsell 1999); a shrinkage of visual receptive fields (RF) and a shift towards the attended location (Ben Hamed et al. 2002; Womelsdorf et al. 2006, 2008); a decreased trial-to-trial variability of individual neuron’s response (Cohen and Maunsell 2009); an increased synaptic efficacy (Briggs et al. 2013); a decrease in noise correlations between neurons (Cohen and Maunsell 2009), and decreased neuronal response latencies (Galashan et al. 2013). This is proposed to have as overall effect to enhance perceptual processing, possibly through local (Chalk et al. 2010; Panagiotaropoulos et al. 2012) and long-range (Popov et al. 2017) neuronal coupling mechanisms.

The correlations we describe between overt behavior and prefrontal target-related spatial representations could be interpreted as a change in the strength of the percept associated with the target or the distractor rather than as a change in the estimate of its spatial position; a strong percept at the time of target correlating with higher probability of correct detections and a strong percept at the time of distractor correlating with higher probability of false alarms. Two arguments speak against this. First, the classification we are applying is not discriminating perception vs. failed perception trials, but rather associating the observed neuronal activities to a spatial estimate. Second, and most importantly, the position of attention orientation in space, as inferred from FEF population activity just prior to target or distractor presentation, is highly predictive of both behavioral speed and accuracy (Monosov and Thompson 2009; Astrand et al. 2016) but also, as we show here, of the spatial estimate of perception. A target has a higher probability of eliciting correct detections when attention is decoded close to this target. Likewise, a distractor has a higher probability of eliciting false alarms when attention is decoded close to this distractor. In other words, both the prefrontal spatial estimates of spatial locus of attention and of target location (as decoded from the population activity) are predictive of behavior.

In conclusion, we show that the population neuronal responses in the FEF not only inform on whether a stimulus has been perceived or not, but also on how accurately it was localized in space, irrespectively of whether the stimulus was an actual target of behavior or an irrelevant stimulus. The accuracy of this spatial representation strongly correlates with overt behavior in terms of speed and accuracy. A strong prediction of this is that in a cued-target detection task in which a spatial response is required (e.g. saccade or pointing), overt error will correlate with the internal prefrontal representation of target location. From a fundamental perspective, while perception is often viewed as an all or nothing variable, our study associates evidence for a measure of reliability of the percept: when a stimulus is detected, this detection can be associated with a very good spatial estimate or with a poor spatial estimate. This view challenges classical models of decision-making or at least calls for the integration of this spatial dimension. From an applied perspective, understanding the neuronal population substrates of stimulus selection, distractor filtering and overt behavior is crucial for developing novel technological advances to improve abilities related to visual discrimination and selection of relevant information in noisy environments or in pathological conditions.

## Acknowledgements

We thank research engineer Serge Pinède for technical support and Jean-Luc Charieau and Fabrice Hérant for animal care. All procedures were approved by the local animal care committee (C2EA42-13-02-0401-01) in compliance with the European Community Council, Directive 2010/63/UE on Animal Care.

